# Analysis of sparse animal social networks

**DOI:** 10.1101/2024.10.31.621436

**Authors:** Helen K Mylne, Jackie Abell, Colin M Beale, Lauren JN Brent, Jakob Bro-Jørgensen, Kate E Evans, Jordan DA Hart, Dabwiso Sakala, Twakundine Simpamba, David Youldon, Daniel W Franks

**Affiliations:** Department of Biology, University of York, York, UK; Leverhulme Centre for Anthropocene Biodiversity, University of York, York, UK; Centre for Agroecology, Water and Resilience, Coventry University, Coventry, UK; York Environmental Sustainability Institute, University of York, York, UK; Centre for Research in Animal Behaviour, University of Exeter, Exeter, UK; Mammalian Behaviour and Evolution Group, Department of Evolution, Ecology and Behaviour, University of Liverpool, Liverpool, UK; Elephants for Africa, London, UK; Gothenburg Global Biodiversity Centre, Gothenburg, Sweden; Department of Geography, University of Zurich, Zürich, Switzerland; African Lion and Environmental Research Trust, Livingstone, Zambia; Department of National Parks & Wildlife, Zambia

## Abstract

Low-density social networks can be common in animal societies, even among species generally considered to be highly social. Social network analysis is commonly used to analyse animal societal structure, but edge weight (strength of association between two individuals) estimation methods designed for dense networks can produce biased measures when applied to low-density networks. Frequentist methods suffer when data availability is low, because they contain an inherent flat prior that will accept any possible edge weight value, and contain no uncertainty in their output. Bayesian methods can accept alternative priors, so can provide more reliable edge weights that include a measure of uncertainty, but they can only reduce bias when sensible prior values are selected. Currently, neither accounts for zero-inflation, so they produce edge weight estimates biased towards stronger associations than the true social network, which can be seen through diagnostic plots of data quality against output estimate. We address this by adding zero-inflation to the model, and demonstrate the process using group-based data from a population of male African savannah elephants. We show that the Bayesian approach performs better than the frequentist to reduce the bias caused by these problems, though the Bayesian requires careful consideration of the priors. We recommend the use of a Bayesian framework, but with a conditional prior that allows the modelling of zero-inflation. This reflects the fact that edge weight derivation is a two-step process: i) probability of ever interacting, and ii) frequency of interaction for those who do. Additional conditional priors could be added where the biology requires it, for example in a society with strong community structure, such as female elephants in which kin structure would create additional levels of social clustering. Although this approach was inspired by reducing bias observed in sparse networks, it could have value for networks of all densities.

## 1 Introduction

Social networks underlie all animal social behaviour, and analysing the factors that affect their structure can improve our understanding of how individuals interact. Social network density forms an important part of social structure, defined by the proportion of dyads (all of the possible pairs of individuals) who could potentially associate within a population that do associate. Some animal social networks are dense, characterised by frequent interactions and connections among a high proportion of its members. Others are sparse, with a high proportion of dyads never associating with one another. This difference in network density can influence an individual’s rate of social learning [1–3] or their risk of pathogenic infection [4–7], which can cascade to population-level impacts. Differences in network density may be due to inherent differences in gregariousness between species, or inter-population variation in resource availability or predation risk imposing different limits on the propensity of different populations to aggregate.

Studies of animal social networks tend to address gregarious species, typically characterised by dense social networks. However, being gregarious and living in groups is not the same as having strong social preferences: two species can be equally gregarious, but one has individuals associating randomly, while in the other they only associate with a few select partners. As of 2019, 78% of published studies focused on the social structure of either birds (e.g., [8–10]) or group-living mammals [11], such as cetaceans (e.g., [12–15]), primates (e.g., [16–18]) and social carnivores (e.g., [19–21]). Understandably, researchers generally focus on highly gregarious species because their associations are more frequent and they are more likely to be co-located so they are easier to sample for social network studies, and there can be clearer applications for the work concerning conservation (e.g., [19]) or captive animal welfare [22]. However, this means that we know only relatively little about other taxonomic groups and species that may have sparser networks, such as solitary mammals (e.g., [23]) or reptiles (e.g., [24]). Less gregarious animals can still have strong friendships or specific preferences regarding who they should associate with, which might impact their behaviour and survival, including species such as African savannah elephants (*Loxodonta africana*), or mountain and western gorillas (*Gorilla beringei beringei* and *Gorilla gorilla gorilla*) that are generally considered highly social but can show relatively low average rates of social association [25–29].

The two core components of a social network are the individuals within the population, called nodes, and the associations between them, known as the edges. In some networks, we only need to know if a connection exists between a dyad, but the strength of that association (henceforth “edge weight”) is not important. In others, we need to know the probability of a dyad being observed together. For many studies, edge weights are the basis of further network structural analyses, making it critical to calculate them reliably.

Edge weight estimation methods are sensitive to the density of a network, so may be problematic when networks are sparse, as we will demonstrate in this paper. Methods designed for dense networks may produce biased measures of edge weight, which can lead to further bias in downstream calculations of node centrality or network density, but this is not something that has previously been examined. Similarly, connections between dyads that do not associate need to be modelled as zero-strength edges, but these methods may be either poor at detecting non-association or falsely identify zero-strength edges. In turn, these errors lead to over- and underestimation of edge weight, which will affect all network studies, regardless of whether they are interested in edge presence or weight. The pitfalls of different methods, which we will identify in this paper, should be considered to decide if zero-inflation is likely to influence edge weight estimation and if edges are more likely to be over-or underestimated.

Animal social network studies often struggle with limited data quality, exacerbating the difficulties of analysing sparse networks using common methods. Frequentist models — such as the commonly-used simple ratio index (SRI [30]) — are biased towards extreme edge weights (0 and 1, respectively indicating that it is impossible for a pair to ever be together or apart), especially when sample size is low [31, 32], and so accept zero edge weight values too readily. They are therefore liable to underestimating network density by overinflating the number of dyads showing non-association. Furthermore, they contain no measures of uncertainty so cannot highlight the dyads whose edge weight is most likely to have suffered from low sample size. In contrast, Bayesian edge weight models can, as with any Bayesian analysis, accept a non-flat prior, making them more reliable when performed correctly. However, they introduce bias if the analyst chooses to use inappropriate priors designed for the analysis of dense networks. These edge weight priors can imply that non-association is highly implausible, producing posterior distributions biased towards stronger associations. This, in turn, can lead to errors in estimating network density. It is, therefore, inappropriate to use either the SRI or a Bayesian model with a prior designed for a dense network when analysing networks that are not fully connected, as we will later demonstrate. So far, this does not appear to have been an issue in publications, as Bayesian edge weight models are a relatively recent innovation and have only been used for populations with dense networks (e.g., [33–40]). However, as Bayesian models generally become more popular, it is likely that researchers will start to use them to calculate the edge weights of less gregarious species, for which a more robust analysis method needs to be established. We can overcome some of the issues with methods designed for dense networks by having very large quantities of data [32, 41], but this is rarely a plausible solution in animal social network studies, and it does not solve all of the issues associated with edge weight estimation, as we will here discuss.

We need a model that reflects how the edge weights are generated in the natural system and incorporates the likelihood of obtaining a true zero, to avoid modelling a connection where one does not exist or biasing our model by supplying inappropriate information regarding the probability of extreme edge weights. All common methods, whether they are frequentist such as SRI (e.g. [10, 21, 24, 42, 43]), half-weight index (HWI) (e.g., [44–47]; or HWIG when correcting for individual gregariousness [48]) or twice-weight index (TWI) (e.g., [49, 50]), or Bayesian such as the Bayesian framework for Inference of Social Networks (BISoN [41, 51]; e.g., [34, 38–40]) or STRAND [52, 53], model a network based on the assumption that the social processes that produce the edge weight contain only a single step: they implicitly assume a fully-connected network where every individual forms a connection with every other, and the only step in deriving the edge weight is determining how strong that connection will be. Edge weight may be correlated with dyad characteristics such as age (e.g., [49]), kinship status (e.g., [25]), phenotypic similarity (e.g., [54]), or anything else that has the potential to induce a preference (or not) for associating, but the decision over how much time to spend together is only a single step.

In contrast to these modelling assumptions, a network that is not fully connected has an additional data-generating step: before defining the weight of the connection, there is a precursory step that determines whether or not a dyad connects at all. This could be driven by a choice to avoid another individual [24, 55] or by spatial or temporal barriers that preclude individuals ever having the opportunity to associate (e.g., one member of the population dies before another is born, or no movement corridor exists between two subpopulations so their home ranges cannot overlap). This creates an additional step that defines the level of zero-inflation in the edge weights. Only once it has been defined that a dyad associates can the edge weight be determined. The sparser a network is, the more important it is to capture the first step of the underlying functional process. It is important to note that while this initial step of edge weight calculation has a Bernoulli outcome, the final network density is not a binary result of dense or sparse but a continuum composed of many dyadic edge weights that lie between zero and one. Models that assume a fully connected network become progressively worse as network density declines, but there is no intrinsic cost to always using a method that captures both steps of the underlying social process.

In this paper we examine two different network modelling approaches for their applicability to sparse networks: the SRI [30], which is the approach that has historically been the most commonly used to estimate the association between individuals, and BISoN [41, 51], which is a newer method of using Bayesian models to estimate the association index with uncertainty and incorporate more appropriate prior information. We demonstrate the application of both of these methods on a sparse network of male savannah elephants. We show how Bayesian models overcome many of the limitations of the SRI but can suffer problems if supplied with priors that are not appropriate for the particular use case. Finally, we introduce a modification of the Bayesian framework that follows a two-step data-generating process, which can deal with the pitfalls of sparse network analysis. We propose that this modification be applied to networks of any density and would benefit future animal network studies.

## 2 Challenges of sparse social networks for current common methods

### 2.1 Common network analysis methods

With gambit-of-the-group data [31], the SRI and Bayesian methods estimate the edge weight between individuals based on the total number of observations in which a dyad was together versus apart. However, despite using the same input data to estimate the same target measure, they suffer from different problems. As with any frequentist measure, SRI estimates are drawn based on the assumption that all possible edge weight values are equally likely — the equivalent of assuming a flat prior — so can produce any output edge weight, even when data are insufficient to support extreme values. To combat this, researchers often only include individuals with total sightings exceeding a threshold value, but taking a subset of the data creates its own issues, discussed below. As a Bayesian alternative, BISoN [41, 51] and other Bayesian approaches such as STRAND [52, 53] can use non-flat priors to inform the model of the prior probability of extreme values, so we can set up the model to be more uncertain about extreme edge values when the dyad observation count is low. Bayesian approaches, therefore, work better with lower sample sizes and uneven sampling than the SRI approach (see supporting information: The SRI is too trusting of extreme values). However, as with the SRI, a single-step prior is appropriate only for fully connected, or close to fully connected, networks. For more sparse networks, we should appropriately modify the prior to reflect the biological processes: here, we consider the underlying two-step social process that generates connections (or not) and then determines the weight of that connection.

The SRI is typically calculated as the number of times a pair of individuals were observed associating divided by the total number of sightings that either individual was observed:

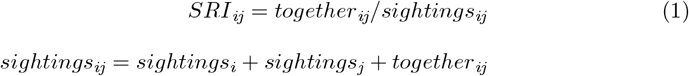

Where *together*_*ij*_ is the number of observations in which individuals *i* and *j* were in the same group; *sightings*_*i*_ and *sightings*_*j*_ indicate the respective number of observations of individuals *i* and *j* in the absence of the other; and *sightings*_*ij*_ is the total number of times they were seen as a whole. As a simple proportion, this measure expects extreme values such as 0 and 1 to be as likely as anything in between. It is a common misconception that frequentist models are less biased than Bayesian models because prior information is not incorporated. This has the opposite effect: a frequentist model still has a prior, but it is forced to be flat and uninformative, so it biases the results towards extreme values, making outliers just as plausible as the average value [56]. The SRI also contains no measure of uncertainty: with this method, a value of SRI = 0 occurs when a dyad is never observed in the same group, with no greater uncertainty around edge weight estimates for poorly sampled dyads than those with many observations, but we should be more certain when more data are available (Fig S1.1).

These issues are the case regardless of network density, but their potential impact on analyses is exacerbated when dealing with extreme edge weight scores. In the real world, a lack of connection between individuals has important implications for social structure, information and disease transfer, and individual centrality. As the true network density declines, the likelihood of obtaining an edge weight of zero (spurious or correct) increases, and the SRI’s lack of uncertainty and treatment of all possible edge weights as equally plausible become a greater issue. We must, therefore, be extremely careful about modelling zero-inflation, but the flat prior and lack of uncertainty seen in the SRI combine to make it very trusting of extreme edge weight values. It is important to note that while we are focussing on the SRI for this paper, other frequentist edge weight measures, including the half-weight and twice-weight indexes, all suffer the same pitfalls. Furthermore, to avoid confusion, it is also worth noting that pre-network permutation tests, which have known problems [57–59], do nothing to deal with the issues faced by frequentist measures.

With Bayesian approaches, we still estimate the edge weight based on the proportion of total dyad sightings in which they were together, but now as a model parameter rather than a simple ratio calculation, for example:

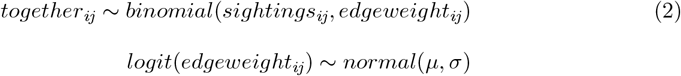

Where *together*_*ij*_ and *sightings*_*ij*_ are the respective number of observations of a dyad in the same group and in total, as above; and *µ* and *σ* are the prior mean and standard deviation. Estimates of edge weight are now described by posterior distributions, which narrow as *sightings*_*ij*_ increases, capturing uncertainty in the edge weight. Incorporating a user-defined prior can limit misleading outputs when the sample sizes are small, as the range of likely edge weight values is reduced. Through this use of sensible priors and quantification of uncertainty, Bayesian approaches are inherently more capable of dealing with the small sample sizes that are common in studies of animal social systems. While BISoN is not the only option available for the Bayesian estimation of edge weight (see also STRAND [52, 53]), we will focus on it in this paper.

### 2.2 SRI requires deletion of data

With a flat prior and no implicit uncertainty measure, limited data availability about a dyad can produce unreliable SRI values, so researchers will usually set an arbitrary minimum threshold number of observations per individual to include that node in the analysis (e.g., [25, 29, 42, 44, 46, 49]). While this does prevent inaccurate edge weights created by the most poorly sampled dyads, it simultaneously introduces new issues [60, 61] that are often overlooked. Firstly, data are often hard-won and already limited, yet this practice requires the discarding of data, sometimes in large quantities. Secondly, data removal may not be completely-at-random [62], which is known to introduce bias [61, 63]. For example, individuals that are less neophobic, more philopatric, or possess distinctive identifying features are more likely to be included in the analysis, because the probability of their having repeat observations is greater. If a study next performs a regression analysis on calculated network measures, this non-random data deletion can directly bias the results, especially if these factors are linked to the exposure or outcome variables in question. Thirdly, the specific value of these observation thresholds often varies between studies, as the actual choice of how many sightings is considered “enough” usually depends on the number of individuals that will remain after filtering out those poorly sampled. For example, Chiyo *et al*. [25] included all male elephants with at least 15 observations in their three-year study period, while Murphy *et al*. Murphy2020-yk used an inclusion threshold of just five observations per four year sampling period. Therefore, comparing the results of different studies and identifying how many sightings are truly “enough” is difficult. Even with several sightings per individual, scores of 0 and 1 can be overinflated relative to their true underlying network, because there are only a certain number of possible edge weight values when sighting counts are low (Fig S1.1).

In contrast, employing a Bayesian framework prevents the requirement for data inclusion thresholds, which improves data quality and reduces biases caused by the possibility of data missing not-completely-at-random.

### 2.3 Bayesian models facilitate the use of more appropriate prior distributions

A model prior should always be carefully considered, which is not possible under the frequentist framework. Given how few papers have thus far used a Bayesian method for edge weight estimation, most social network analyses are based on flat frequentist priors, which are generally inappropriate (if nothing is known at all of the population in question, then a flat prior may still be a reasonable choice, though this would be extremely unusual). The *bisonR* [51] package for running BISoN models supplies very broad default priors, but the package authors strongly recommend tailoring them to each specific scenario. Most social networks contain at least some true zeros. Any model, frequentist or Bayesian, that uses a flat or very broad prior will struggle to model these zero values because they are modelling only the second step of what is inherently a two-step process.

Unlike the SRI, with BISoN we can change our model structure to incorporate two steps — first to determine if a dyad will associate at all, and second to define the strength of that connection — if we supply it with an appropriate prior, which package default priors are not. Here, we have presented a prior allowing BISoN to model both steps without changing the social network analysis workflow. While we could instead use a mixture model with zero-inflation to model a two-step process, using either a frequentist (see supporting information: Standard zero-inflated model) or Bayesian framework, we can keep the overall method relatively simple by changing only the prior structure. All network models should include some option to account for zero-inflation, and selecting the best method for measuring edge weights requires careful consideration: Bayesian techniques require integration of current understanding of the system, and a prior predictive check, which can lead to biases if carried out using inappropriate priors, but their ability to estimate uncertainty and accept non-flat priors makes them more appropriate than frequentist methods.

### 2.4 Prior selection in a Bayesian edge weight model

Bayesian methods have been successfully applied to dense networks, using both simulated and empirical data [34, 64, 65], and their use in primate research has been discussed in subsequent publications [66, 67]. However, the best way to use them for sparse networks has not yet been evaluated. Prior choices require careful consideration for any analysis, not just social networks. Fig 1a shows an improper uniform prior, in which all edge weights are considered equally likely before exposure to the data. By relaxing the assumption that all edge weights are equally probable, we can make the prior more informative. When we demonstrate the use of BISoN on an example dataset, we first show the outcome of this uniform prior, followed by the default prior supplied in the *bisonR* package [51]: a wide symmetrical prior (logit(*edge weight*_*ij*_) *∼ normal*(0, 2.5)) that presumes very little about the average edge weight (Fig 1b). Third, we show the result of using a strong asymmetrical prior (*edge weight*_*ij*_ *∼ beta*(1, 5)) that indicates weaker edges are the most likely but still allows for some strong associations (Fig 1c). How we define the prior depends on our knowledge and assumptions of the system: the sparser the network, the more right-skewed the prior distribution should be.

**Fig 1.**
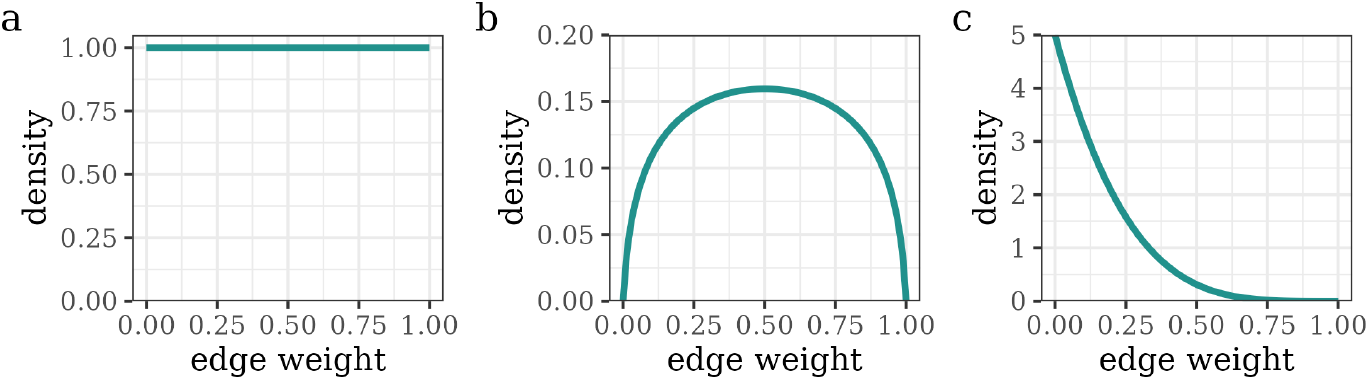
Prior assumptions for single-step BISoN models. Prior assumptions for a) SRI or a BISoN model with a uniform prior, b) a BISoN model with a weak, symmetric prior (logit normal, *µ* = 0, *σ* = 2.5) and c) a BISoN model with a strong asymmetric prior (beta, *a* = 1, *b* = 5).

Biases can appear in the edge posterior when the prior distribution does not reflect the underlying social process: knowing that the network is of a lower density can allow us to adapt our prior to a more suitable distribution, and the assumptions we can make based on that knowledge are critical to dealing with poorly sampled dyads. When the prior average is a long way from that of most dyads, it induces a systematic shift in the direction the data pushes the posterior from the prior. In a sparse network, many of the dyads will never associate, so a prior designed for a dense network will usually be shifted towards peaking at zero by the data. The data for poorly sampled dyads will, by definition, not be able to shift the posterior as far from the prior as the data from the well-sampled dyads, so if they all shift in the same direction, the level of sampling could potentially have a greater effect on the final edge weight than the actual raw data values. When this shift is consistently towards zero, poorly sampled dyads, on average, receive edge weight distributions with higher average values than well-sampled dyads. It is difficult to imagine a scenario in which the number of observations of a dyad would be strongly dependent on the strength of their social bond or vice versa, therefore this correlation highlights a problem with the edge weight data. We show this happening in the example analysis. Again, without well-sampled data, the choice of prior (including the SRI’s uniform prior) becomes critical to the success of the edge weight estimation and all subsequent analyses.

## 3 Introducing a two-step prior for sparse networks

By varying the prior distribution, Bayesian methods can be tailored to a sparser or denser network. The modification that we propose simply extends this ability: to alter the BISoN prior structure to reflect a two-step social process producing the edge weights, in which step one defines the extent of zero-inflation and step two the weight of the non-zero social bonds. We structure our model to use two separate edge weight priors, selecting which to use depending on whether a dyad has ever been observed together (Fig 2), such as:

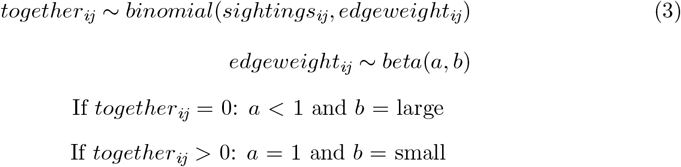

**Fig 2.**
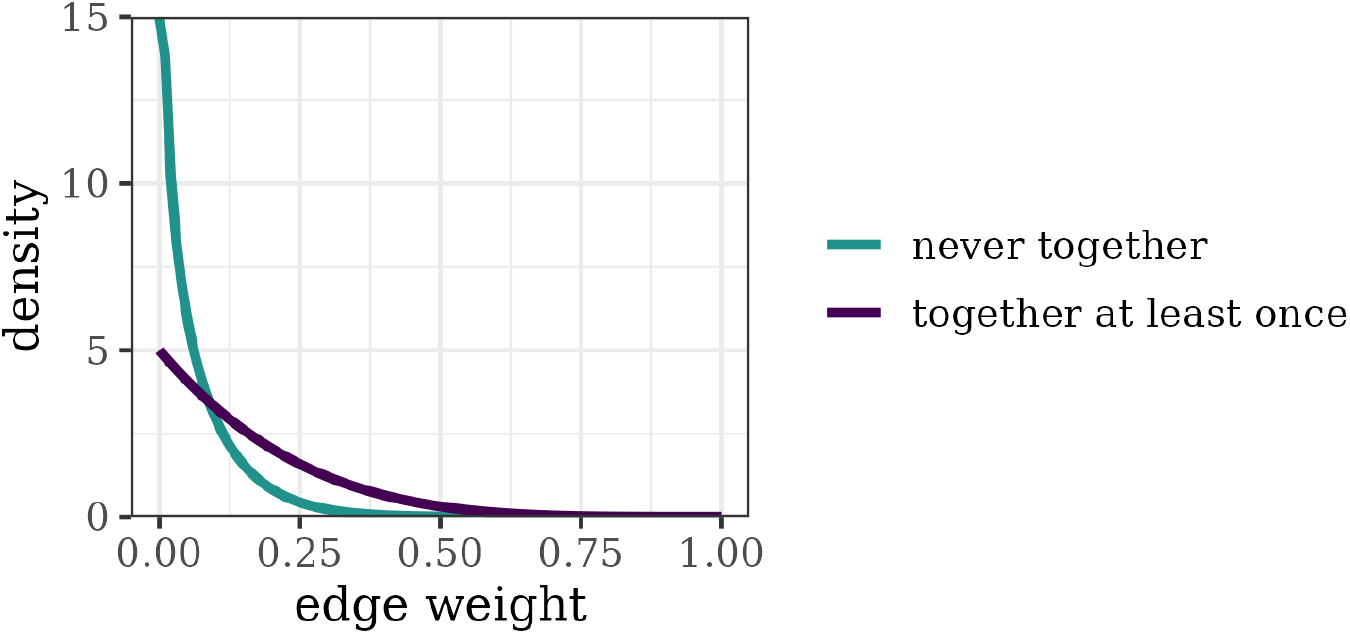
Conditional edge weight prior for a BISoN model of a sparse network. The blue line indicates the lower prior that will be selected when a dyad has never been observed in the same group (in this case *beta*(*a* = 0.7, *b* = 10)). The purple line shows the upper prior choice for when a dyad has been observed grouping together on at least one occasion (in this case *beta*(*a* = 1, *b* = 5)). While in this model we have chosen to still have this prior peak at zero, this reflects that we still expect very low edge weights amongst those that do associate, and the data immediately overwhelm the possibility of zero being an option. Note that the upper prior is the same as the strongly right-skewed prior shown above.

Where again *together*_*ij*_ refers to the number of times a pair was observed in the same group, and *sightings*_*ij*_ is the number of times they were observed in total. This is not a major change to the method itself, as BISoN has always been capable of using a conditional prior. The change is that by capturing both parts of the underlying social process in a conditional prior, we only need to change one part of our overall method.

The new prior is simple to apply: if a dyad is never observed in the same social group, then the model will use a prior that increases the probability of an edge weight of zero; if a dyad has been observed interacting on at least one occasion, it will receive an edge weight calculated using a prior with less or no zero-inflation. This, therefore, allows the model to identify the most appropriate prior and incorporate the probability of non-association as an initial step before identifying the best edge weight distribution per dyad. As before, poorly sampled dyads will return an edge weight more similar to their respective prior, while well-sampled dyads will shift further. The model becomes less uncertain about a zero edge as the total sighting count increases: the uncertainty in these values indicates that zero is the most likely edge weight but still allows stronger relationships as a possibility when sampling is poor. In contrast, for a dyad observed together on at least one occasion, we use a wider prior that allows higher edge weights to be observed, again becoming more confident in the results as total sampling increases.

The most logical step here may appear to be to make our regular prior more right-skewed so it is closer to zero. However, if we were to use only an extremely right-skewed prior, such as the lower prior of this combination, to encapsulate the zero-inflation and the true associations, a prohibitively large amount of association data would be required for any truly strong edge weights to register.

The values a researcher might choose for the prior depend on their study species and expected edge weights. In the following example, we have opted to have the prior for associated dyads peaking at zero, but this is immediately overwhelmed by the data. This produces edge weight values that are still far more likely to be close to zero than to one, based on the evidence that male elephants do not appear to form strong relationships with one another [25, 26, 29]. If we were expecting to see a strongly bimodal distribution in the final edge weights, with some dyads never associating and others showing consistent association, we would shift our upper prior further from zero.

## 4 Example dataset

For this example, we used data on sparse networks based on associations between male elephants of the Mosi-Oa-Tunya National Park (MOTNP), Zambia, collected from May 2016 until December 2017 (all permissions for data collection were supplied by Department of National Parks and Wildlife, Zambia: permit number TJ/DNPW/101/13/18). No additional ethics permission was required alongside the research permit, but to avoid causing any stress to the elephants, we monitored them from a vehicle from a safe distance, and kept all noise to a minimum while in the National Park.

This population is relatively small, totalling approximately 500 individuals, with similar numbers of males and females. The park is well connected to large areas of suitable habitat, and the elephants are free-ranging, so individuals may leave the area for long periods. Data were collected during daily drives through the park, marking down the identities of all individuals. From these observations, we created 504 days of gambit-of-the-group [31] data comprising 213 males sighted in 481 groups. Finally, we converted this to a data frame containing every possible dyad with counts of the total number of times that dyad had been observed and how many of those occasions they were in the same group. Data analyses were run by calling Stan (version 2.26.1 [68])from R (version 4.2.1 [69]) using the *cmdstanr* [70] package.

## 5 Results from empirical data

### 5.1 Simple Ratio Index (SRI)

When we calculate the SRI values for this population (Fig 3a), we get an extremely high proportion of zeros (17385, equating to 77.00% of dyads). While a much lower number of dyads receive an SRI of one (12 dyads, equating to just 0.05% of the total population), the likelihood of these apparent permanent alliances would be exceedingly small given the population average.

**Fig 3.**
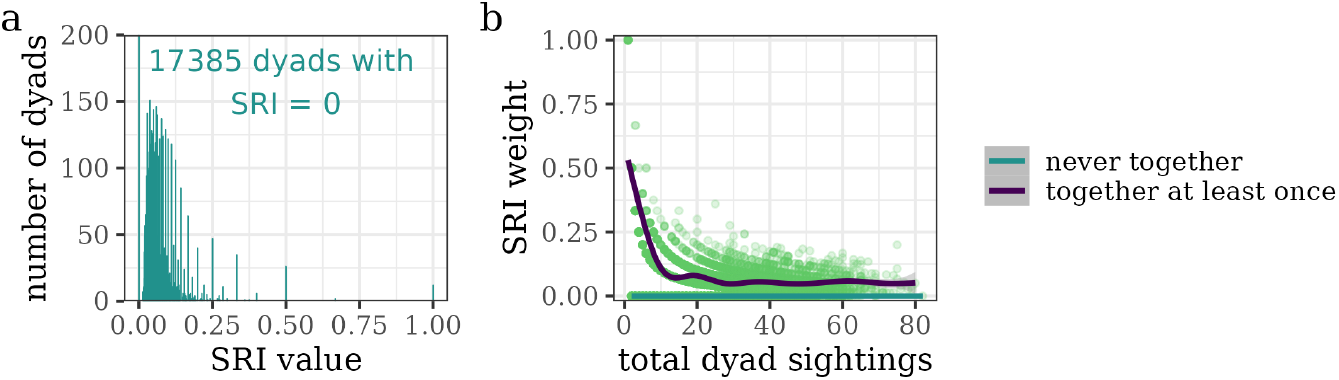
Edge weights for MOTNP elephants using the SRI. a) The vast majority of dyads have an SRI score (green points) of zero, indicating that the output is certain that these pairs never associate with one another, regardless of the number of sightings. b) This translates to a large split between the edge weights calculated for poorly sampled dyads that have (purple line) or have not (blue line) ever been observed grouping together. We see a very strong effect of dyad sighting count on the edge weight for dyads observed together on at least one occasion, until we reach around 12 sightings per dyad.

Given this, the standard next step with the SRI would be to set a threshold for the minimum number of sightings per node required to be included in the analysis. If we set this threshold to five sightings per node, the proportion of dyads with an SRI score of zero shifts down to 46.13%, and all dyads scoring a 100% association rate are removed. A threshold of 10 sightings per individual drops the 0% association rate down to 26.25%, showing high instability in the SRI outputs. This is highlighted by Fig 3b, which shows the association between the total number of observations per dyad, and the calculated edge weight for that dyad, stemming from the extreme number of zeros calculated by the SRI. The overall results would strongly depend on the threshold value we choose, which, as previously stated, is an arbitrary choice.

We could potentially use Fig 3b to determine a less arbitrary inclusion threshold by selecting the point at which the patterns in the edge weights weaken. However, this example shows that this is somewhere between 10 and 20 sightings per dyad. This would require removing at least all individuals with fewer than 10 observations. In this case, that would take our total sample size down from 213 to just 74 elephants. To remove 65% of the sample could severely bias our results if the remaining portion is not entirely representative of the total population. Therefore, we cannot rely on objective means to determine the threshold level.

### 5.2 Bayesian model with a single-step prior

When we use the BISoN framework with a single-step prior, we can see a limited effect of prior shape on the posterior distribution, but the inclusion of zero as a possible edge value is very important. When using a fully flat or highly right-skewed prior, in which zero is a possible option, we see very similar posterior distributions (Fig 4a and c), whereas for a normal prior peaking at 0.5 for which extreme edge weights are unlikely (and actual values of 0 and 1 are impossible) we see an overall much broader and less skewed posterior (Fig 4b).

**Fig 4.**
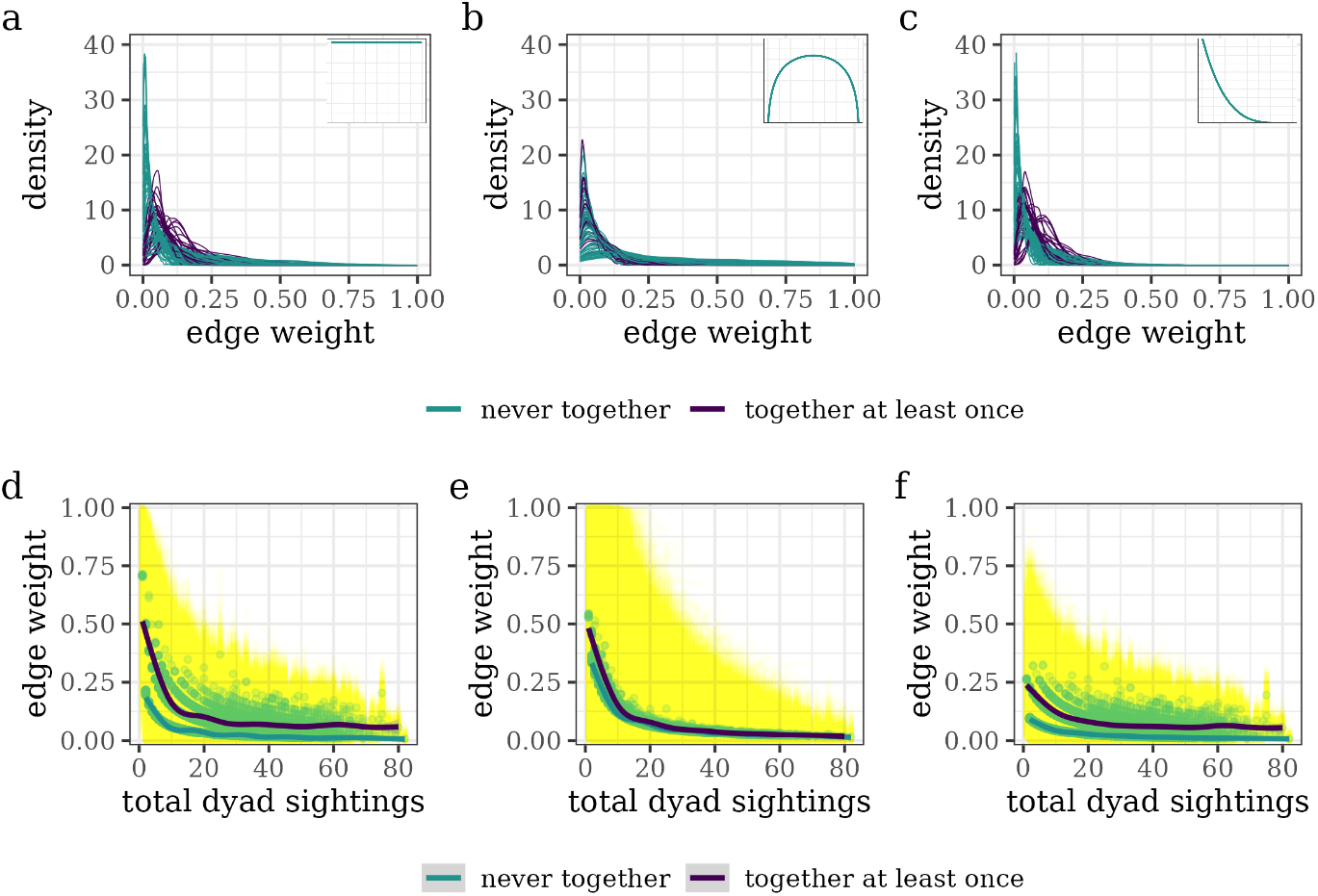
Edge weights for MOTNP elephants using BISoN models with single-step priors. Results using BISoN with uniform (left), default (middle) and right-skewed (right) priors (insets in top row show the respective prior for the column). a-c) The top row shows the posterior distribution for 100 randomly sampled dyads, with green curves indicating pairs that have never been observed in the same group and purple curves showing dyads that were observed in the same group on at least one occasion. The right-skewed and uniform priors produce posterior distributions that initially appear very similar because they still allow an edge weight of zero to be a plausible draw, whereas the default prior draws all values together. d-f) The bottom row shows the effect of total sighting count per dyad on their median (green points) and full (yellow points) posterior distribution. The blue line indicates dyads never observed together, while the purple line shows dyads observed together at least once. As the prior becomes increasingly right skewed, the effect of sighting counts on median edge weight declines.

We now also have a situation in which there is a negative correlation between mean edge weight and total observations per dyad at low sighting counts, especially when using very broad priors. Looking at Figs 4d and 4e, we can see that poorly sampled dyads appear to show higher edge weights than well-sampled ones (an effect the authors have also seen in social network data for macaques). Again, this is most severe for the logit normal prior, which draws all of the values towards the middle instead of creating variation between dyads. The effect is also stronger for the flat prior than for the right-skewed, for which the trend is almost entirely removed because those dyads with the fewest sightings do not have sufficient data to discredit high edge weight model draws, so the posterior is very wide, with high median values. Similar reasoning also explains the differences between BISoN with a flat prior and the SRI: with the SRI, low sighting counts largely score exactly zero, whereas with BISoN low sighting counts are given a very wide posterior (shown in yellow) so their median (green) is substantially above zero.

Based on these graphs, the best solution would appear to be to use BISoN with a strongly right-skewed prior, which can model the edge weights without creating strong patterns of social association based on data quality. However, this prior may not allow sufficient movement towards a higher edge value for dyads showing a strong social bond. Therefore, we need a prior that better reflects the underlying social process, such that it allows for some dyads to attain higher edge weights, but without increasing the prior average and causing poorly sampled dyads to consistently obtain a higher edge weight than well-sampled ones. Our modification to BISoN to use a prior that reflects the two-step underlying social process facilitates edge weight estimation in a manner robust to zero inflation and which allows stronger edges to be estimated.

### 5.3 Bayesian model with a two-step prior

When using the two-step prior, the posterior distributions of the edge weights (Fig 5a) appear very similar to those with single-step priors, though with greater confidence in some of the weakest edges, as we would expect. The median edge weight is associated much more weakly with the total number of sightings than with the broader priors (Fig 5b), even at low dyad sighting counts. While the shape of the curve for pairs seen grouping together is very similar to the single-step right-skewed prior, the overall effect of low sighting count has been removed by the model’s increased ability to obtain median estimates very close to zero. We now have edge estimates that are more reliable than the previous BISoN models, as they use a prior that allows the modelling of zero-inflation.

**Fig 5.**
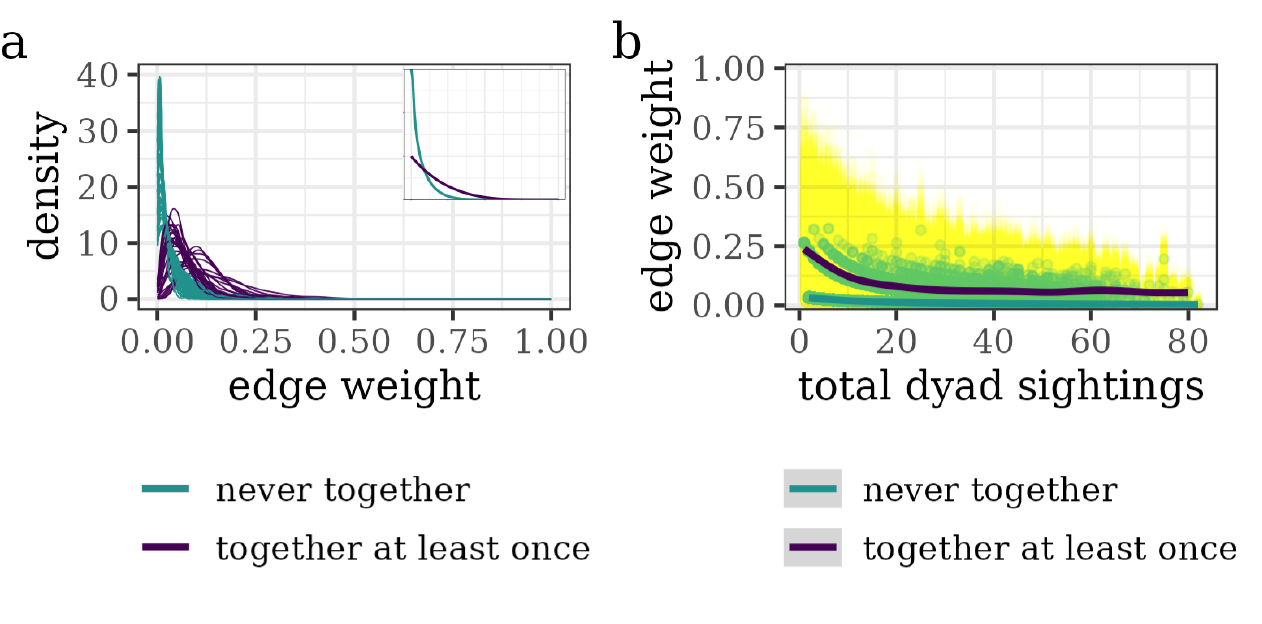
Edge weights for MOTNP elephants using a BISoN model with a conditional prior. a) posterior density lines for every dyad (blue indicates density lines drawn for pairs never observed in the same group, purple for those seen together on at least one occasion); and b) edge weight as a response to number of sightings per dyad (full posterior distribution in yellow, median per dyad in green). While some effect of sighting count on the median edge weight for individual dyads remains, the impact is much reduced compared with the previous BISoN plots, and it lacks the values of 100% association observed with the SRI.

### 5.4 Using downstream network measures to assess edge weight quality

In general, we should be able to see directly in our edge weight values if there is a problem in our estimation method. Sometimes however, the impacts of the method are masked until subsequent analyses of the outputs. For example, in an analysis to identify the effects of individual trait values on network centrality, we may find that our centrality values contain spurious patterns, such as in the number of sightings or number of different partners that an individual is observed with compared to network centrality. Here, we will show that using an inappropriate single-step prior for edge weight can severely impact downstream analyses of network structure. We explore this using eigenvector centrality, a measure of how connected an individual is to other well-connected individuals, as it is commonly used and is the centrality measure most robust to causal analysis [71]. As with the edge weights, we can plot measures of centrality calculated per model against a) the number of other individuals in the population with whom they were never observed associating, and b) their individual sighting count, and look for any unexpected trends that may indicate problems with the outputs.

When we plot the SRI-based eigenvector data against the number of non-associated elephants, we see a negative correlation (Fig 6a), exactly as we would expect. If, however, we compare the SRI-based centrality to the number of sightings per individual, we see a positive correlation for poorly sampled dyads (Fig 6b), which indicates that there is a problem with the estimation. This relationship is created by the extreme edge values caused by limited numbers of samples, especially for elephants observed fewer than 10 times. This could be used as an objective method of determining an inclusion threshold, but as before, this would require the discarding of more than half the data. We obtain similar trends when calculating the edge weight using a zero-inflated mixture model (Fig S2.2).

**Fig 6.**
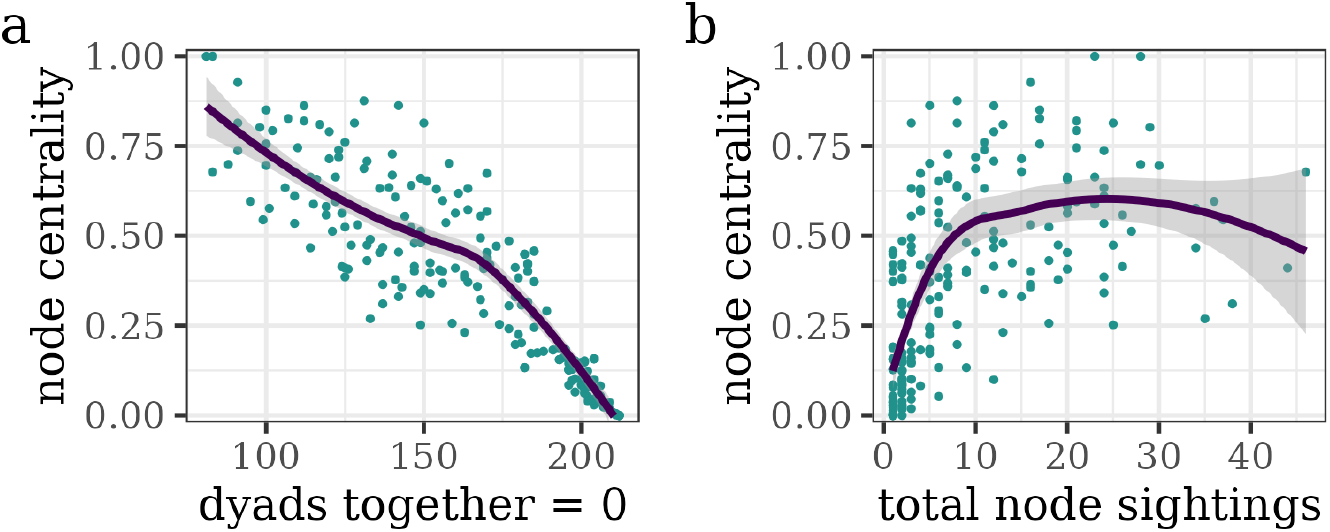
Eigenvector centrality estimates based on the SRI edge weight calculations. a) There is a negative relationship between SRI-based centrality and the total number of potential partners with whom a focal was never observed associating, which is as expected. However, b) indicates an issue with the SRI-based centrality in that there is a strongly positive effect of the number of observations per individual on their social position at low observation counts.

We see something very different when we look at the eigenvector centrality plots based on the single-step BISoN models. Startlingly, there is an apparent reversal of the effect of the number of non-associated individuals in the population, such that elephants observed with the most partners receive the lowest centrality scores, while those observed with the fewest partners receive the highest (Fig 7a-c). This is, of course, nonsense. Eigenvector centrality as a measure combines the number of association partners an individual has with the number of association partners their partners have. The most central elephants therefore cannot be those with the fewest association partners. The edge weight estimation process must have an underlying error to create this trend.

**Fig 7.**
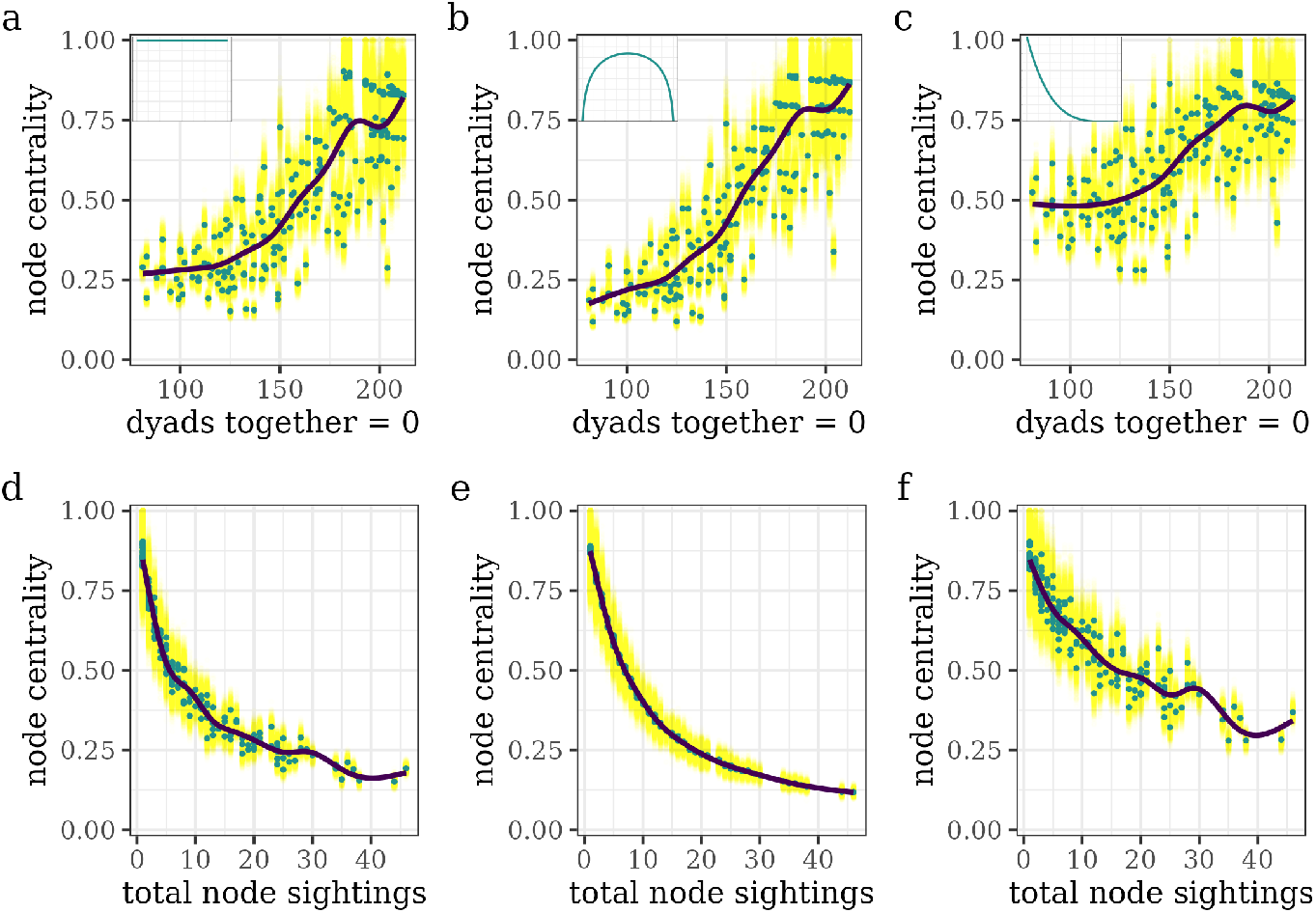
Eigenvector centrality estimates based on BISoN edge weight models with single-step priors. Priors per column are shown as insets on the top row: left to right uniform, default and right-skewed. In all graphs, yellow points indicate the uncertainty around the median, displayed in blue. a-c) The top row indicates the impact of number of dyad partners with whom an individual was never observed associating, where we see a spurious positive effect. The more right-skewed prior suffers more weakly from this than the uniform-based or default-based centrality, but still sufficiently to create serious cause for concern. There is no biologically or statistically meaningful explanation for this to be a correct trend. The pattern is explained by d-f) in which the elephants with the most observations occupy the social positions of lowest centrality.

This seemingly impossible pattern occurs because of an underlying trend in the number of observations per elephant (Fig 7d-f). Individuals with fewer overall observations contribute fewer sightings to dyad observations. When the edge weight prior is designed for dense networks, the posterior for most dyads is shifted closer to zero from the prior. As the data for poorly sampled dyads are less influential on the distance the posterior can move, dyads with fewer observations consistently appear to have a stronger edge weight than well-sampled dyads. When this translates through to individual effects, and poorly sampled individuals end up in more poorly sampled dyads, it is exaggerated so individuals observed rarely have the strongest average edge weights and, therefore, the highest eigenvector centrality.

Finally, we can now consider eigenvector centrality calculated from the edge weights produced using BISoN with a two-step prior and see that we successfully avoid all downfalls from the other methods. Changing to a conditional prior has successfully reversed the spurious correlation between number of associates and node centrality (Fig 8a) by removing the effect of sighting count on eigenvector (Fig 8b). We have avoided creating estimates biased by the data quality, so any subsequent analyses using these data will be more reliable than those calculated using an improper prior while also avoiding the need to remove poorly sampled dyads as we would with the SRI.

**Fig 8.**
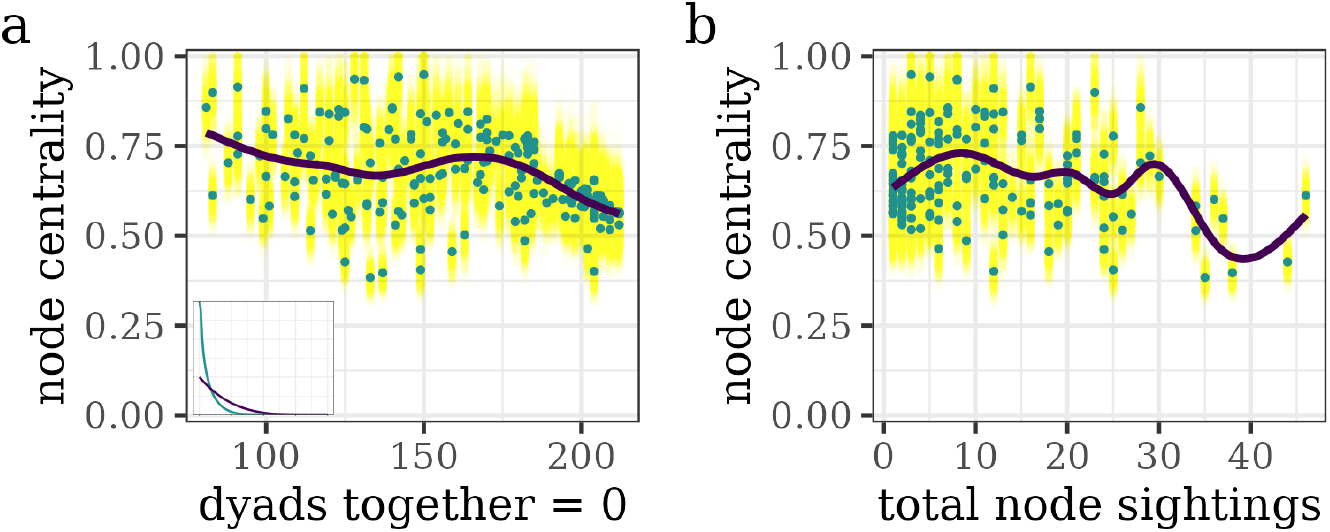
Eigenvector centrality estimates based on BISoN edge weight model with a two-step prior. a) Using our modified prior (inset), to reflect a two-step social process, we have successfully restored the negative trend in node centrality such that those seen with the most different dyad partners on average have the highest centrality, and those with the fewest have the lowest. b) Individual sighting count now has extremely limited effect on eigenvector centrality.

This quick sanity check of the eigenvector against the number of associates can instantly highlight if there is a problem with the edge weights, and show how sighting count could affect centrality. Our modification does not entirely eliminate this, and care should be taken to look for any anomalies. For example, in the MOTNP data, there is a single sighting where 89 elephants crossed the river and came up into the surrounding mopane forest together, of which 31 were males with known identities. For two males, this was their only observation. Therefore, these two individuals were “always” seen with one another and are in the “seen grouping together at least once” group for 30 dyad pairs. This means that, despite having very wide uncertainty margins for all dyads due to low total counts, they will still have an above-average median edge weight due to the nature of their single observation. When carried forward to a centrality analysis, these elephants will display a slightly elevated eigenvector centrality. Using the modified BISoN framework means this elevation is much lower than when using an unconditional prior or the SRI, and the number of individuals for which this could be an issue is reduced, limiting the impact on the regression. However, we still need to be aware of the potential for data artefacts that an occurrence such as this may leave. It is worth noting that while none of these methods is perfect, by using a method that incorporates current knowledge, quantifies uncertainty, and does not induce issues in downstream analytical data frames, we can rest assured that we are using the best possible option.

## 6 Discussion

We have shown that current animal social network analysis methods do not cope well with sparse network data. Frequentist measures such as the SRI suffer from an innate and unavoidable reliance on a flat prior, which overestimates the confidence that can be placed on extreme edge weight values. Combine this prior with an inability to incorporate uncertainty, and researchers must rely on setting arbitrary inclusion thresholds when analysing poorly sampled dyads, leading to data loss and potentially biased results. While Bayesian models can account for these problems in dense networks, they can still produce edge weights overly influenced by the frequency of sightings per dyad when inappropriate priors are applied to sparse networks. By modifying the BISoN framework to use a conditional prior, reflecting a two-step underlying social process, we can create a model where the edge weights are more reliable than those from the SRI, a zero-inflated mixture model, or an uninformed Bayesian model. As such, they are no longer overly dependent on data quality, which is commonly poor in animal social network studies.

When checking our models, we used several diagnostics to compare output quality and determine if our BISoN modification improved the current methods. The first question is whether a result fits with what we know of the population: the SRI produces far too many dyads with zero probability of ever associating, even when we set a high inclusion threshold. At just 66 km^2^ [72], the MOTNP is not sufficiently large to preclude the interaction of elephants by simply never meeting, with a bull easily able to cross it in a single day. It is, therefore, extremely unlikely that this many dyads are truly never together, as there would not be enough space for them all to move without encountering other elephants. This allows us to make two informed assumptions: i) the proportion of zeros produced by the SRI is overinflated; and ii) that well-sampled dyads that are never recorded associating have elected not to do so, rather than happening to never encounter one another. When using BISoN, the conditional prior can help to alleviate the effect of sighting count on median edge weight by using the broader prior with a higher average only when we have evidence that the dyad has elected to sometimes associate. While this solution does not entirely remove the impact of limited sightings on edge weight, it does minimise it compared to a single-step prior.

At first glance at the edge weights alone, it seems that the conditional prior is not particularly advantageous over the right-skewed single-step prior: the basic outputs for the edge weight look fairly similar. However, we have modelled the zero-inflation, which, while it appears to have only had a limited effect on the edge weights themselves, has a very noticeable and important impact on our subsequent analyses. The node centrality values produced can act as a suitable sanity check. While there appears to be little difference between the right-skewed BISoN outputs and the two-step BISoN outputs when we consider only the edge weights, the eigenvector centrality from a single-step prior in a sparse network creates an impossible situation in which individuals with more partners receive a lower centrality score. This trend is corrected by the use of a two-step prior. We can use such checks to ensure our outputs make sense in downstream analyses.

Selecting the two distributions remains important for successful edge weight estimation when applying a two-step prior to a sparse network model. Making the zero-centred prior too strong will not allow enough variation when data are limited, but if too weak some of the possible edge weights may still present as too high and create trends in edge weight and centrality based on sighting counts. Similarly, if the probability mass for the non-zero prior is not shifted far enough from zero, any stronger relationships will be missed; shift it too much, and the same trends of increased edge weight with reduced sightings will be observed within the group who have been observed together. In this instance, we have opted for a large overlap between the priors such that dyads only seen once or twice will receive relatively similar posterior distributions regardless of which prior they are assigned to. By giving both priors a peak at zero, a dyad observed just once but who were together at that time will still have a low average edge weight (though the zero-peak is always immediately overcome by the data), matching our assumptions of a sparse network. This particular conditional prior is the one that works for our specific elephant population and is not a silver bullet prior that will work for all network analyses.

Every author should identify their own priors to create a set that works for their own population, which does not have to require that both priors peak at zero: unlike with the male elephants, a network can be sparse but still contain some strong associations, in which case an upper prior that allows higher edge weights may be preferable. For example, if we were to repeat this analysis with the female elephants of MOTNP, we would expect a more bimodal distribution in our output because of the fission-fusion structure of female elephant society [73–75]. In this case, we could shift our non-zero prior to be more distinct from the lower, producing a bimodal distribution of edge weights, indicating strong social bonds within certain dyads. It could even be taken a step further to add more prior distributions as necessary. Again, using the example of the female elephants, we may, in this scenario, want the prior to reflect a three-step social process: a zero-centred prior for those never seen in the same social group, a right-skewed or more central prior for those who have been seen grouping together but who are unrelated; and a left-skewed prior for related females. Hierarchical networks in which we see kinship groups within subgroups, which themselves join together to form larger association groups are very common in animal societies. This method could be easily expanded to allow for the use of different priors depending on which level of association a dyad is expected to fall into.

It should be noted that the assessment of the edge weight is rarely the end of the analysis. Generally, we calculate the edge weights so that we can later assess trends in node-, dyad-or network-level characteristics. Therefore, the edge weight calculation only creates the data ready for a future regression model. There is no issue with checking the edge weights produced — just as we would check for errors in any other raw data — and running the model again using an improved prior if there is a problem. If edge weight outputs are dependent on the dyad sighting count, or there are issues with downstream calculations such as the centrality, there is no issue with adjusting the prior, so long as it still reflects our original understanding of the population.

In conclusion, the shape of the prior is, as with all analyses, the key to estimating edge weight, and using a prior structure that reflects the underlying social process is especially important for sparse networks, though the concept applies to dense networks too. With a frequentist model, this means considering the implications of a flat prior and including in our discussion any impacts this may have on the results. For a Bayesian model, it means tailoring the prior to our individual situation rather than using the broad default priors supplied by statistical packages, which will usually allow too much probability density over highly improbable edge weight values. In the case of sparse networks, whether the model is Bayesian or frequentist, we need to model the zero-inflation induced by the first step of the underlying social process. While a zero-inflated mixture model can do this better than the SRI, we still recommend that a Bayesian model with a conditional prior is a better option. This means allowing edge weights to sample down to zero using a lower prior and the possibility for stronger relationships with an upper prior. When investigating denser networks, our prior distributions may differ, but we can still consider the underlying social process as containing more than one step, even if the reduced proportion of zero relationships masks the effect. We can therefore use a conditional prior in a Bayesian framework to identify greater nuance in social network data, for networks of any density.

## 7 Acknowledgements

We would like to thank the African Lion and Environmental Research Trust (ALERT) for providing the example dataset and the Zambian Department of National Parks and Wildlife (DNPW) for allowing its collection and analysis. Analyses for this paper were run on the Viking cluster, a high-performance computing facility provided by the University of York. We are grateful for computational support from the University of York, IT Services and the Research IT team.

---

**S1 File. The SRI is too trusting of extreme values**. Simulation plot to show the difference in potential outputs from SRI versus a Bayesian estimator given true values and the number of times a pair were observed.

**S2 File. Standard zero-inflated model**. Frequentist two-step model outputs as an alternative option to BISoN.

