## Supporting Material for "Analysis of sparse animal social networks"

### Supporting information

**S1 File**

#### S1 The SRI is too trusting of extreme values

One of the problems with the SRI is that it is highly sensitive to the amount of data available per individual and dyad, yet it fails to account for this during edge weight estimation. Fig S1.1 shows a simulated example, in which the true dyad edge weight is highlighted in yellow. The purple bars indicate the variation of possible values that the SRI could theoretically take. For example, for a dyad only seen twice, the possible number of sightings they have together are 0, 1 or 2, so the theoretical SRI values are 0, 0.5 or 1. For a dyad seen 10 times, the possible number of sightings together is any integer from 0-10, meaning the possible SRI values are 0, 0.1, 0.2 and so on through to 0.9 or 1. It is not possible with only 2 sightings to obtain an SRI of 0.7, or with 10 sightings an SRI of 0.35. In Fig S1.1, the height of the purple bars indicates the probability of obtaining each of these theoretical values, based on the probability of observing each combination of sightings together versus apart given the true proportion of time spent together.

In all cases, the most likely SRI value is indeed the one that matches the true edge weight. When a dyad are truly together 10% of the time, and they are observed on 10 occasions, the most likely combination of sightings is nine apart and one together, giving an SRI of 0.1. However, the combined probability of all other bars (i.e., the probability that a pair spending 10% of their time together are seen together during a different percentage of their observations) indicates that the probability of accurately calculating the edge weight using SRI is relatively low.

This elevated probability of incorrect estimates over correct ones is due to the SRI working off of the assumption of a fully flat prior, in which all possible values are equally likely. This therefore makes it too willing to accept extreme values, and the

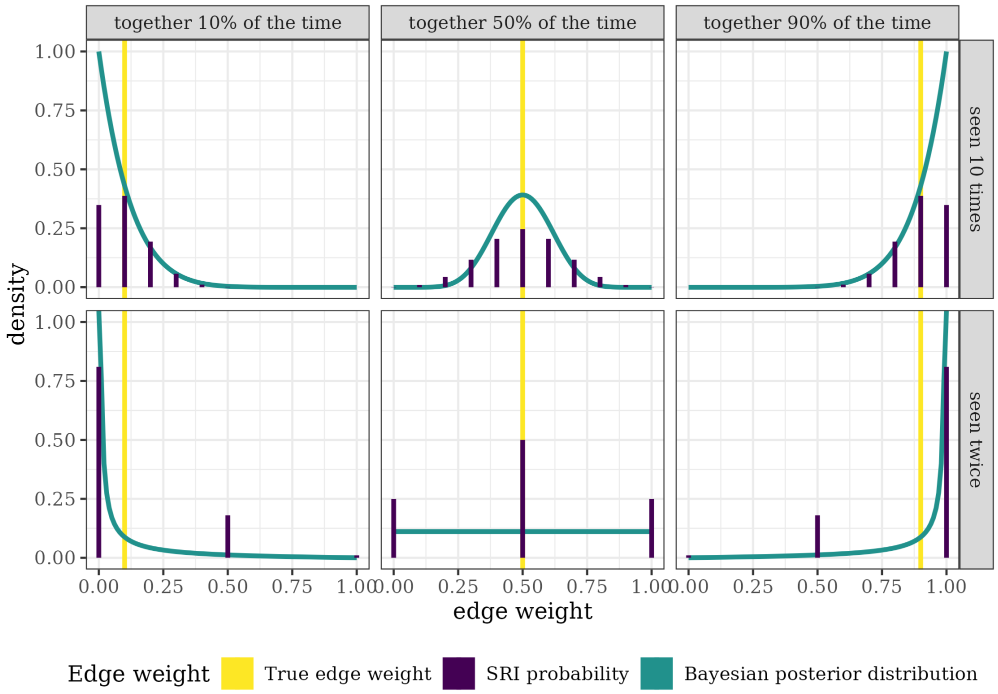


**Fig S1.1. Simulated SRI and Bayesian output values.** Possible outputs for a hypothetical SRI and Bayesian edge weight analysis. In each panel, the yellow line indicates the true simulated value for the length of time that a dyad spends together. The purple lines indicate the possible values of the SRI edge weights, with the height showing the probability of obtaining that value based on the probability of obtaining different combinations of together versus apart during sampling. Only one SRI value will be derived per dyad, showing that the probability of inaccuracy is extremely high. Blue lines show a Bayesian posterior distribution. A Bayesian model measures the association rate with uncertainty, allowing us to be more confident about well-sampled dyads than poorly-sampled ones, and providing an indication of the full distribution of plausible edge weights. Furthermore, that distribution is tailored to match our prior beliefs of the system, so there is less chance of severe misestimation caused by an inherently flat frequentist prior.

researcher is forced to work with purely the observed data, regardless of whether this reflects what is already known. This is further exacerbated by the issue that, as a single point-estimate per edge, the SRI displays no uncertainty, despite the potential for the estimate to be far from the true value, and after the values have been calculated there is no way to tell apart a dyad with an edge weight of 0.5 from 10 or 100 sightings and a dyad with an edge weight of 0.5 from just two sightings.

**S2 File**

#### S2 Standard zero-inflated model

To model a two step process does not necessitate that we use a Bayesian framework. The choice to still use BISoN over a frequentist model comes down to the multitude of reasons outlined in this paper regarding the problems with flat priors and the ability to incorporate uncertainty into edge weight estimates. However, if we were to choose a frequentist framework for this model, we could still model a two step social process with the same data by using a mixture model instead of the SRI.

Mixture models allow the use of a zero inflation term to model the first step of the edge weight derivation process. Here we used the package glmmTMB [1] to create a binomial model with zero inflation of the form:

$$cbind(together_{ij}, apart_{ij}) \sim (1|dyadID_{ij}) \tag{1}$$
$$apart_{ij} = sightings_i + sightings_j$$

Where $together_{ij}$ is the number of observations in which individuals $i$ and $j$ were in the same group, and $sightings_i$ and $sightings_j$ indicate the respective number of observations of individuals $i$ and $j$ in the absence of the other.

This model produces an output that has similarities to both the SRI and the conditional BISoN model. As in the SRI, the outputs are frequentist. They are therefore single-point estimates, so have no inherent uncertainty calculated within them, and are based on a prior assumption that all possible values are equally likely. However, the inclusion of the zero inflation term has now produced an overall bimodal distribution (Fig S2.1a) that more accurately reflects the assumptions of the conditional BISoN.

The model tells us that the global logit edge weight is very low at -4.378 $\pm$ 0.0228, but with a large variance among dyads of 1.094. When we plot the edge weight against the number of sightings per dyad, we see no effect at all of sighting count on the edge weight, but the edges are all measured as extremely weak. On the face of it, this looks like it could be the best of all the models, but when we check the eigenvector values for it, we see the same issues as in the SRI: there is a strong positive correlation between number of sightings per node and the eigenvector centrality, and unlike the SRI this extends as high as nearly 30 sightings before reaching a plateau (Fig S2.2). If we were

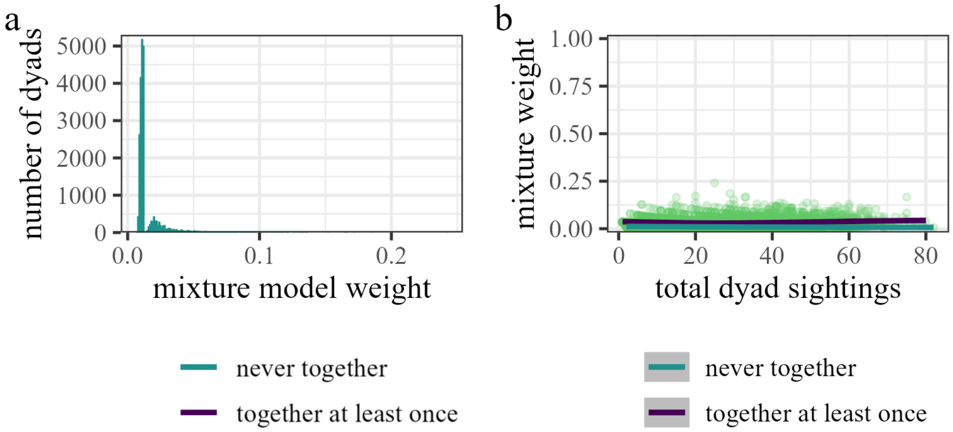


**Fig S2.1. Edge weights calculated using a frequentist zero-inflated mixture model.** a) The distribution of estimates for the whole population is bimodal, indicating the zero inflation and two-step underlying process. The edge weights are much closer to zero than produced using the other methods, creating a situation in which b) the total sightings have very little influence at all on the edge weight calculated, for either dyads that have (purple) or have not (blue) been observed in the same group.

to remove all elephants with fewer than 30 observations from the analysis dataset, we would be left with only nine individuals from our original 213, dropping 96% of our network. Given this, we still recommend using a conditional prior in a Bayesian framework over a frequentist mixture model.

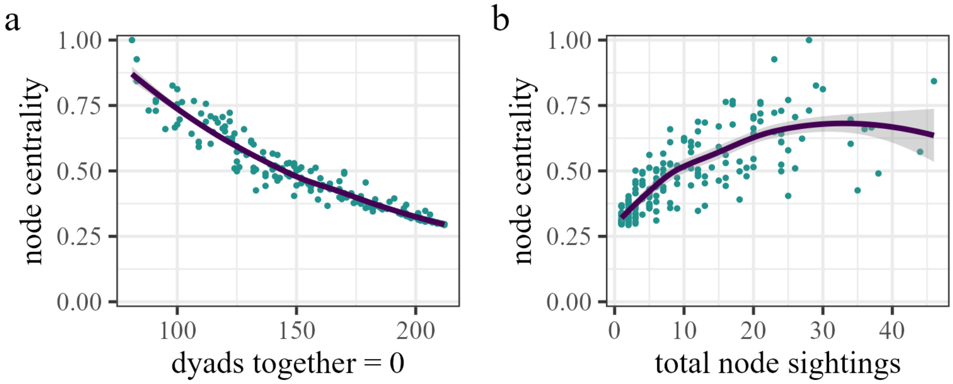


**Fig S2.2. Centrality estimates from a zero-inflated mixture model** a) There is a negative correlation between the number of potential dyad partners that an elephant never associates with and their eigenvector centrality calculated from a binomial zero-inflated mixture model, which is as we would expect. However, as with the SRI, we also see b) a positive effect of total number of sightings of an individual and their node centrality score, which is not a trend that we should observe.
